# ATP-Free Fatty Aldehyde Biosynthesis Enables an Autonomous Lux-Based Bioluminescence System

**DOI:** 10.64898/2026.07.30.741946

**Authors:** Subhan Hadi Kusuma, Takeharu Nagai

## Abstract

Autonomous bioluminescence systems enable continuous light emission in engineered organisms by genetically encoding both luciferase enzymes and their substrate biosynthetic pathways, offering a powerful platform for non-invasive and long-term monitoring of biological processes. However, bioluminescence output is highly sensitive to substrate availability and host metabolic state, often leading to signal instability under energy-limited conditions. Here, we report an alternative luciferin biosynthetic strategy for the bacterial Lux bioluminescence system in which the adenosine triphosphate (ATP)-dependent LuxEC complex is replaced by α-dioxygenase (αDOX), an enzyme that directly converts fatty acids into fatty aldehydes without consuming ATP. Using machine learning–guided directed evolution, we engineered αDOX variants that markedly enhanced bioluminescence intensity when coupled with bacterial luciferase. The resulting ATP-free Lux bioluminescence system enabled single-cell–level bioluminescence imaging and maintained stable light emission under diverse antibiotic treatments, demonstrating enhanced robustness against metabolic perturbations.

**SIGNIFICANCE:** Autonomous bioluminescence imaging has become an attractive method for long-term observation of biological phenomena without the need for exogenous substrate addition. However, the light output of existing autonomous bioluminescence systems, including bacterial and fungal pathways, remains ATP-dependent and often declines when cellular metabolism is perturbed, limiting their reliability for quantitative analysis. Here, we report the development of an ATP-independent substrate biosynthesis pathway for the bacterial luciferase system using αDOX. To improve system performance, we applied machine learning– guided directed evolution, which significantly enhanced signal intensity and enabled single-cell bioluminescence imaging. Furthermore, the αDOX-based bacterial luciferase system maintained stable luminescence under antibiotic treatments, in contrast to the conventional ATP-dependent bacterial luciferase system. In summary, our findings establish a robust ATP-independent autonomous bioluminescence imaging platform that enables monitoring of cellular events under metabolic perturbations.

## INTRODUCTION

Bioluminescence-based reporter systems have been widely used to monitor biological phenomena in living cells^1–3^. Bioluminescent signal generation requires the genetic expression of a luciferase enzyme together with its chemical substrate, luciferin^4^. Conventional bioluminescent reporters, such as firefly luciferase (FLuc) and NanoLuc (NLuc), have been extensively applied for bioluminescence imaging in various host expression systems due to their high signal-to-noise ratios, lack of photobleaching and phototoxicity compared with fluorescent protein biosensors, and wide dynamic range^5–7^. However, these systems depend on the exogenous supply of luciferin, which limits continuous signal acquisition, increases experimental cost, and suffers from poor substrate penetration and uneven distribution in tissues ^8,9^.

To enable sustained bioluminescence without external substrate supplementation, autonomous bioluminescence systems derived from bacterial and fungal organisms have been developed, in which the genes encoding both luciferase and luciferin biosynthesis are genetically integrated^10–12^. In both systems, however, luciferin biosynthesis requires adenosine triphosphate (ATP) (**Figures 1A and S1**). Consequently, fluctuations in cellular metabolic activity—such as those induced by nutrient deprivation or metabolic inhibitors—can lead to reduced luminescence intensity^11^. Importantly, this decrease can occur independently of bioluminescence gene expression driven by inducible promoters, thereby compromising the quantitative accuracy of autonomous bioluminescence systems as reporters and biosensors ^10,13^.

**Figure 1.**
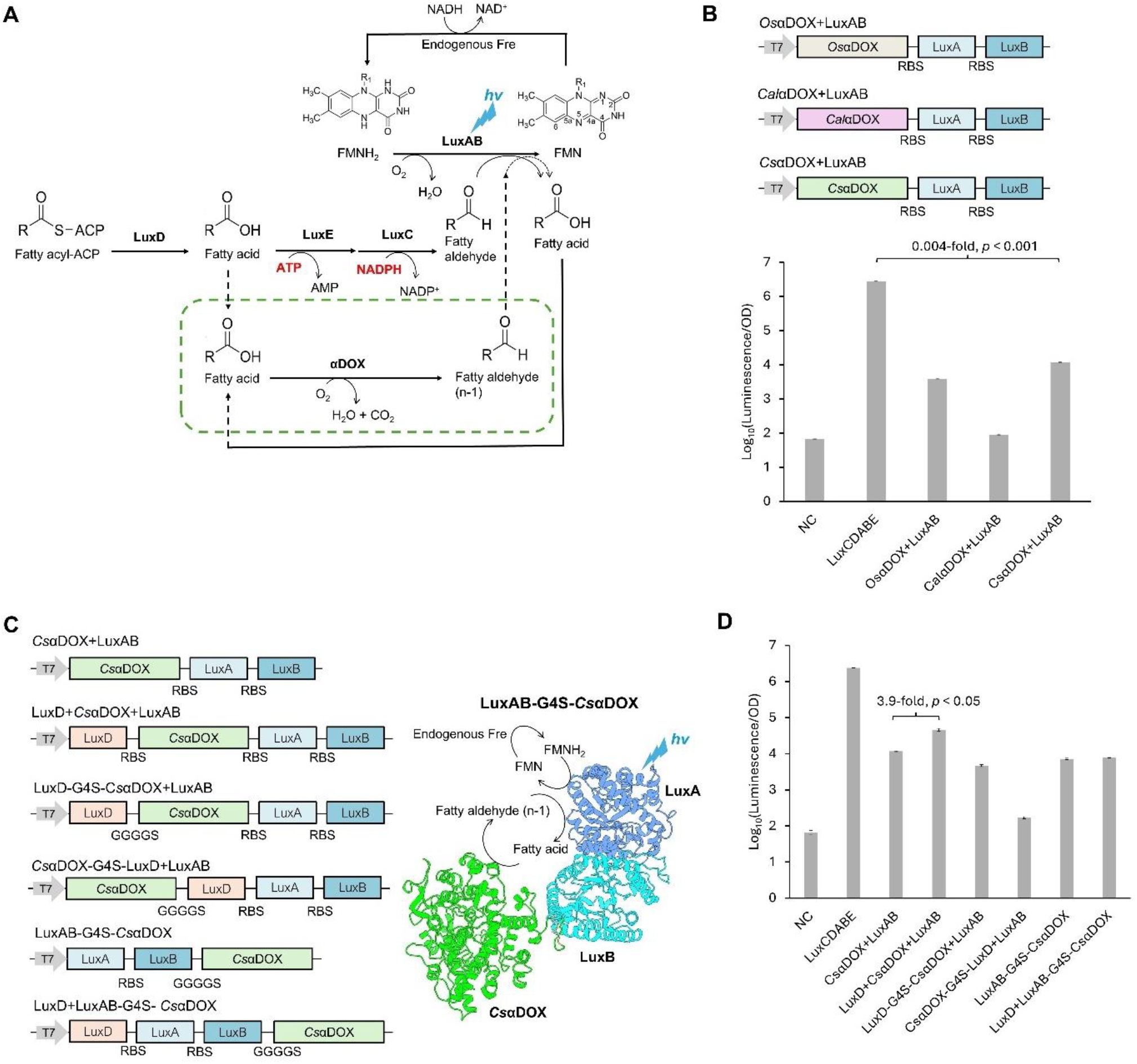
Design of ATP-independent autonomous bioluminescence system. (A) ATP-independent bacterial bioluminescence pathway catalyzed by α-dioxygenase (αDOX). (B) Schematic of αDOX genes optimization (upper panel) and the resulting bioluminescence intensity (bottom). NC, non-transformant *E. coli* JM109(DE3). Data are presented as mean ± SD. *n* =3 technical replicates. (C) Schematic of *Cs*αDOX-Lux operon (left) and structural model of a single autonomous bacterial bioluminescence system (right). (D) Bioluminescence intensity of the optimized *Cs*αDOX-Lux operon variants. NC, non-transformant *E. coli* JM109(DE3). Data are presented as mean ± SD. *n* =3 technical replicates and normalized by OD_600_.

Here, we report on the development of an ATP-free autonomous bioluminescence pathway based on the bacterial Lux system. Specifically, we replaced the native LuxEC complex ^14^, which produces fatty aldehyde luciferin through an ATP-dependent fatty acid activation process, with α-dioxygenase (αDOX)^15,16^. αDOX is an oxygen-dependent enzyme that directly converts fatty acids into fatty aldehydes (n-1) without ATP consumption, using molecular oxygen and producing water and carbon dioxide as byproducts ^15,17^ (**Figure 1A**). To enhance bioluminescence output, we optimized αDOX through machine learning–guided directed evolution, yielding variants that produced substantially increased luminescence when coupled with bacterial luciferase (LuxAB). Furthermore, the resulting αDOX-based Lux bioluminescence pathway maintained stable light emission under multiple antibiotic treatments, whereas luminescence from the native LuxCDABE operon was markedly attenuated under the same conditions. Together, these results establish the αDOX–Lux system as a metabolically robust, energy-independent autonomous bioluminescence platform.

## RESULTS

### Design of an ATP-Independent Autonomous Bioluminescence System

As a starting point, we selected the bacterial luciferase system as a platform for autonomous bioluminescence (auto-bioluminescence) because of its superior solubility and thermostability compared with fungal bioluminescence systems^10,11^. In the native biochemical pathway (**Figure 1A**), luciferin biosynthesis is mediated by a three-enzyme cascade consisting of LuxD (thioesterase), LuxE (acyl-protein synthetase), and LuxC (acyl-CoA reductase)^18^. LuxE and LuxC form a fatty acid reductase complex that catalyzes the production of long-chain fatty aldehydes in an ATP- and NADPH-dependent manner^18,19^.

To establish an alternative long-chain fatty aldehyde biosynthesis pathway that does not require ATP as a cofactor, we investigated α-dioxygenase (αDOX) enzymes^16^. We selected three well-characterized αDOX homologs derived from *Oryza sativa* (*Os*αDOX)^20,21^, *Crocosphaera subtropica* (*Cs*αDOX)^15^, and *Calothrix parietina* (*Cal*αDOX)^22^. These αDOX genes were introduced into a *luxAB* expression system, and all three αDOX-containing systems produced detectable autonomous bioluminescence in *Escherichia coli* JM109(DE3), clearly distinguishable from the non-transformed negative control (**Figure 1B**). Among the tested enzymes, *Cs*αDOX yielded the highest bioluminescence intensity relative to the other αDOX homologs. The result is consistent with previous reports showing that *Cs*αDOX preferentially utilizes medium-chain fatty acid substrates (C10–C16)^15^, which are more compatible with bacterial luciferase activity^13^. In contrast, *Os*αDOX preferentially acts on longer-chain fatty acids (>C16)^15,21^, while *Cal*αDOX exhibited comparatively poor performance, likely due to protein aggregation associated with its intrinsic structural properties^22^. Despite these differences, the luminescence intensities of all αDOX-based systems remained lower than that of the wild-type LuxCDABE operon, indicating that further optimization is required to achieve brightness comparable to the ATP-dependent pathway.

The autonomous bioluminescence intensity of the Lux system is influenced by the intracellular availability of fatty acids, which serve as precursors for fatty aldehyde production. Accordingly, enhancing fatty acid supply—either through overexpression of fatty acid biosynthesis enzymes, such as thioesterases or through exogenous myristic acids supplementation—can modulate the performance of the LuxCDABE and *Cs*αDOX–Lux operons. To test this, we examined whether overexpression of thioesterases, including LuxD, ‘TesA^23,24^, and TesB^25^, could enhance fatty acid biosynthesis in *E. coli* (**Figures 1C and S2A**). Introduction of thioesterases led to increased bioluminescence intensity, with LuxD producing up to a 3.9-fold enhancement compared with ‘TesA and TesB (**Figures 1D and S2A**). These results indicate that fatty acid availability is a rate-limiting factor for fatty aldehyde production in *E. coli* expressing *Cs*αDOX and LuxAB. Consistent with this interpretation, supplementation with myristic acid resulted in a concentration-dependent increase in luminescence intensity, with an optimal concentration of 200 µM. Under these conditions, bioluminescence intensity increased by up to 19-fold relative to *E. coli* cultures without myristic acid supplementation (**Figure S2B**). Furthermore, incorporation of LuxD into the *Cs*αDOX+LuxAB system enabled a 9.4-fold increase in bioluminescence intensity while requiring substantially lower concentrations of myristic acid (50 µM) compared with the *Cs*αDOX+LuxAB system lacking LuxD (**Figure S2C**). These results indicate that LuxD alone is sufficient to enhance fatty acid availability in *E. coli*, thereby supporting higher luminescence output, although further optimization through additional metabolic engineering may be required to maximize performance. In addition, *E. coli* expressing LuxCDABE operon exhibited up to a 4.3-fold increase in luminescence intensity upon supplementation with a low concentration of myristic acid (20 µM) compared with the *Cs*αDOX–Lux system (**Figure S2D**). Under these conditions, *in vivo* fatty acid conversion to fatty aldehyde appears to be more efficient in the LuxEC-dependent pathway than in the *Cs*αDOX-Lux system.

We further investigated a fusion-protein strategy designed to accelerate fatty aldehyde channeling by minimizing diffusional distance between enzymes. In this design, LuxD and LuxB were fused to the N- and C-termini of *Cs*αDOX, respectively. However, this fusion strategy did not lead to an increase in bioluminescence intensity (**Figure 1C and 1D**). Given the relatively large molecular weight of *Cs*αDOX (70.5 kDa)^15^, impaired protein folding or reduced enzymatic activity likely contributed to the limited performance of the fusion constructs. Consistent with this interpretation, detectable luminescence from the LuxD–G4S–*Cs*αDOX fusion was observed only after 48 h of incubation at room temperature, suggesting delayed protein folding or maturation of the fusion protein (**Figure S3**). Despite these limitations, the LuxAB–G4S–*Cs*αDOX fusion architecture offers conceptual insight into the construction of a single-complex autonomous bioluminescence system and may serve as a starting point for future structural or protein-engineering optimization.

### Directed Evolution Guided by Machine Learning Improves αDOX–Lux Bioluminescence Output

To further enhance bioluminescence intensity, we performed directed evolution of *Cs*αDOX using a machine learning–guided strategy. Specifically, we applied protein language model (PLM)–based mutagenesis to improve both the fitness and catalytic activity landscapes of *Cs*αDOX. As an initial step, we employed a PLM-based zero-shot approach to explore the fitness landscape using EvoProtGrad^26^. EvoProtGrad, also referred to as plug-and-play directed evolution (PPDE), integrates unsupervised and supervised models based on a discrete Markov chain Monte Carlo (MCMC) framework ^26^. In subsequent rounds, we focused on enhancing the activity landscape using EVOLVEpro^27^, a method previously shown to improve the catalytic performance of diverse enzymes, including CRISPR nucleases, prime editors, and T7 RNA polymerase. Notably, both EvoProtGrad and EVOLVEpro employ ESM-2^28^ as the underlying PLM for mutational design. Following four iterative rounds of mutagenesis, we generated and evaluated a panel of *Cs*αDOX multi-mutant variants (**Figure 2A**).

**Figure 2.**
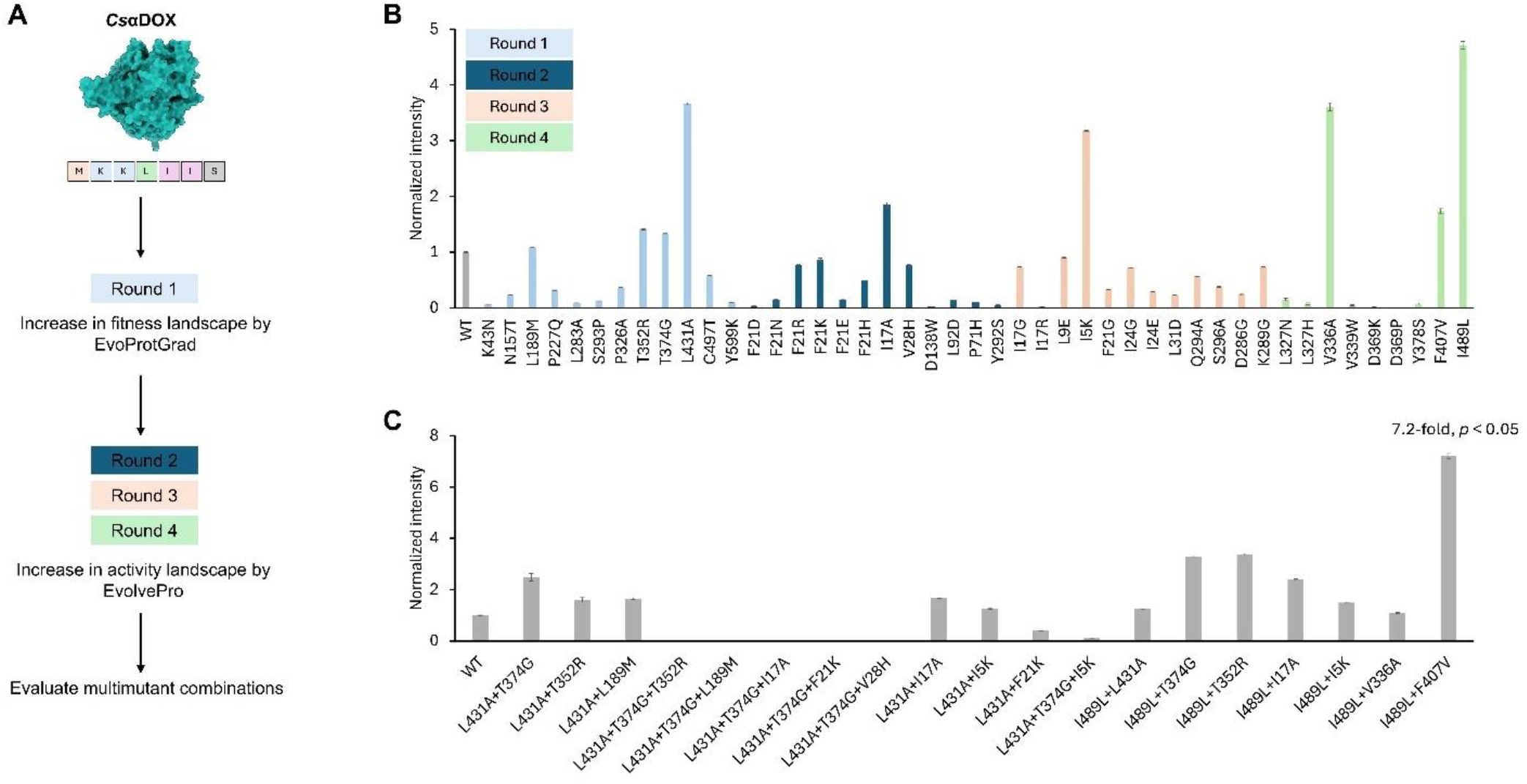
Screening of *Cs*αDOX-Lux Operon libraries by machine learning–guided directed evolution. (A) General scheme for continuous evolution workflow. The evolution cycle was first guided by EvoProtGrad to screen the fitness landscape, followed by EVOLVEpro to progressively enhance the activity landscape, and finally by evaluation of multimutant combinations. (B and C) Results from the first to fourth rounds of *Cs*αDOX evolution (B) and evaluation of multimutant *Cs*αDOX combinations. WT, wild-type *Cs*αDOX-Lux operon. Data are presented as mean ± SD. *n* =3 technical replicates and normalized by OD_600_.

The brightness of the *Cs*αDOX-Lux operon (LuxD+*Cs*αDOX+LuxAB) increased as early as the first round of mutagenesis (**Figure 2B**). Screening of *Cs*αDOX multi-mutant variants identified the *Cs*αDOX-I489L/F407V mutant (hereafter referred to as enhanced *Cs*αDOX, e*Cs*αDOX), which increased bioluminescence intensity by up to 7.2-fold compared with the original *Cs*αDOX–Lux operon (**Figure 2C**). In parallel, and prior to *Cs*αDOX mutagenesis, we performed directed evolution of LuxA, the catalytic subunit of bacterial luciferase, using EvoProtGrad and EVOLVEpro. LuxA mutagenesis was conducted in the context of the native LuxCDABE operon. This screening yielded a brighter LuxA variant harboring the mutations C130E/N148G/F186N/A128F (LuxA v.1), which exhibited approximately 10-fold higher brightness than wild-type LuxA. Introduction of an additional mutation, Q344A (eLuxA) further increased brightness to approximately 14-fold relative to wild-type LuxA (**Figure S4**). Subsequently, LuxA v.1 and eLuxA were incorporated into the e*Cs*αDOX–Lux operon, resulting in up to a 7.9-fold increase in brightness compared with the original *Cs*αDOX–Lux operon (**Figure 3A**). To further improve photon output, we next introduced a yellow fluorescent protein (Venus) to increase the apparent quantum yield via bioluminescence resonance energy transfer (BRET), as previously demonstrated in the development of Nano-lanternX system^12^. Fusion of Venus to eLuxA yielded the e*Cs*αDOX–eYNLX operon; however, this modification did not result in a further increase in brightness compared with the e*Cs*αDOX–eLuxA operon alone (**Figure 3A**).

**Figure 3.**
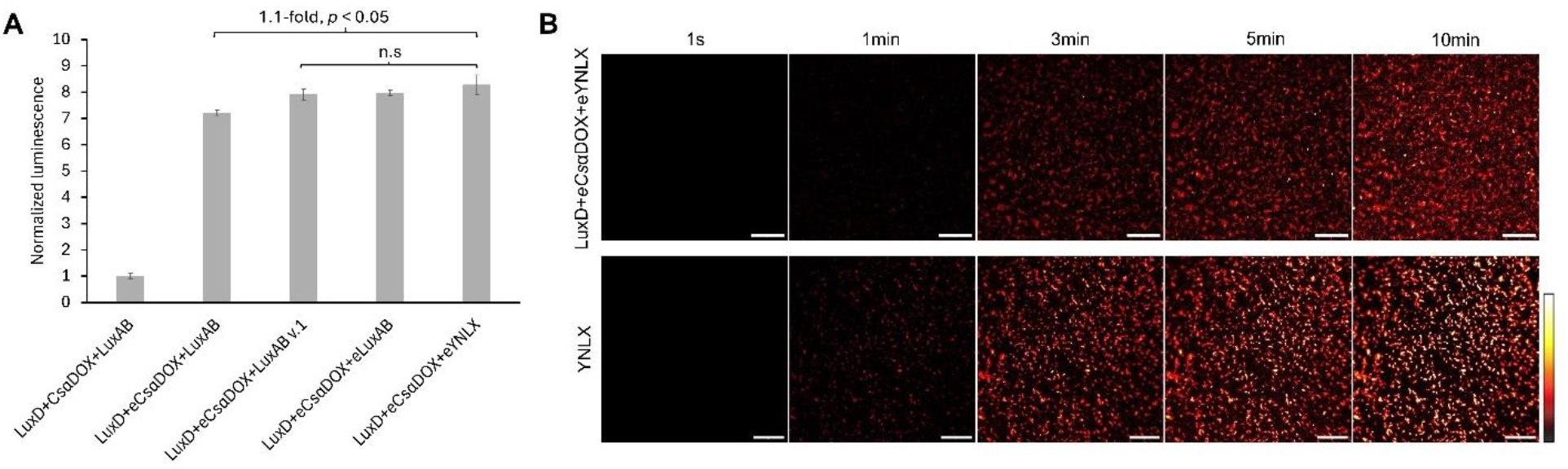
Introducing of enhanced LuxA and bioluminescence imaging of the *Cs*αDOX-Lux operon. (A) Bioluminescence intensity of *Cs*αDOX–Lux operon variants incorporating enhanced LuxA and the yellow fluorescent protein Venus. Data are presented as mean ± SD. *n* =3 technical replicates and normalized by OD_600_. (B) Autonomous bioluminescence imaging of *E. coli* expressing e*Cs*αDOX-eYNLX and YNLX operon. Pseudocolor images, scale bars, 20µm; 60× magnification with indicated exposure times.

### Bioluminescence Imaging and Assay of e*Cs*αDOX-Lux Operon

The e*Cs*αDOX–eYNLX strain was next evaluated for bioluminescence imaging. Bioluminescence signals from e*Cs*αDOX–eYNLX were readily detectable with an exposure time of 3 min. In contrast, the YNLX operon exhibited stronger single-cell bioluminescence signals than the e*Cs*αDOX–eYNLX operon under identical imaging conditions (**Figure 3B**). Notably, the original *Cs*αDOX–Lux operon produced luminescence signals that were too weak to be detected by bioluminescence microscopy (**Figure S5**).

Because the *Cs*αDOX–Lux system is an ATP-independent autonomous bioluminescence system, we next examined whether its luminescence intensity is affected by antibiotics that impair cellular viability. We tested several antibiotics with distinct mechanisms of action, including kanamycin (translation inhibitor)^29^, meropenem (cell wall biosynthesis inhibitor)^30^, and rifampicin (transcription inhibitor)^31^. Lux expression systems were driven either by a constitutive T7 promoter using the original pRSET ^B^ vector (carbenicillin resistance) or by an inducible T7 promoter containing the lac operator (T7*lac*) using a modified pRSET^B^ vector (carbenicillin resistance). For the inducible system, the *lac* repressor gene (*lacI*) was included, and 1% glucose was added to suppress leaky expression. After overnight cultivation, cultures were adjusted to an OD^600^ of 0.5 prior to antibiotic treatment.

In the constitutive expression system (**Figures 4A and 4B**), luminescence from both the LuxCDABE and e*Cs*αDOX-eLuxA operons increased during the initial ~10 min of observation. However, in *E. coli* cells expressing LuxCDABE, however, luminescence declined markedly within 10-20 min following treatment with meropenem or rifampicin and exhibited fluctuating signals with an overall declining trend upon kanamycin treatment. In contrast, cells expressing the e*Cs*αDOX-eLuxA operon maintained sustained luminescence under all antibiotic conditions tested.

**Figure 4.**
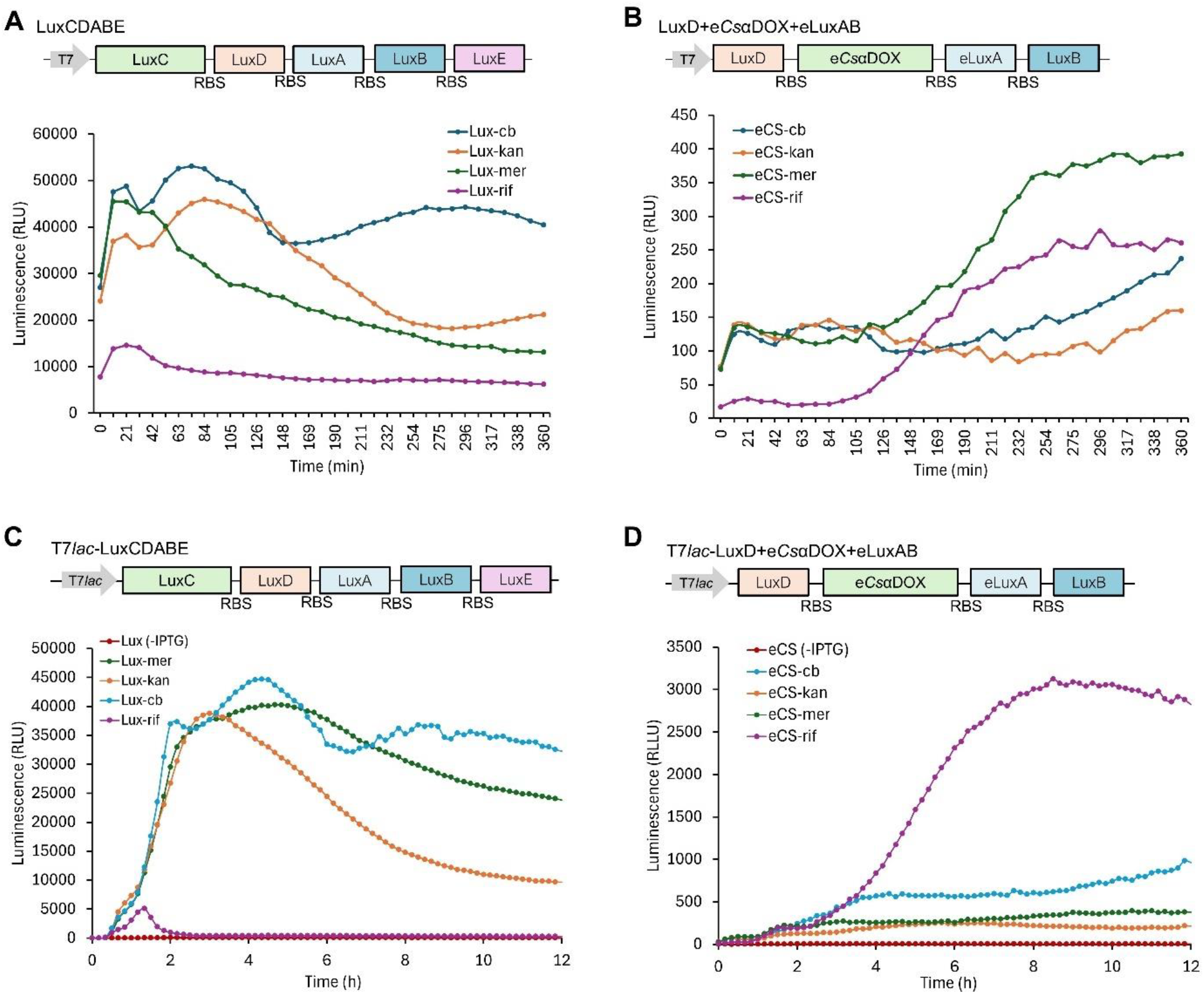
Antibiotic response of *E. coli* JM109(DE3) expressing LuxCDABE and e*Cs*αDOX-eLux operon. (A and B) Bioluminescence dynamic following the addition of different antibiotics in constitutively expressed LuxCDABE (A) and e*Cs*αDOX-Lux operon (B). Data are presented as mean ± SD. *n* =3 technical replicates and adjusted to OD_600_ = 0.5. (B and C) Bioluminescence dynamics following antibiotic treatment in inducible expressed LuxCDABE (C) and e*Cs*αDOX-Lux (D). Data are presented as mean ± SD. *n* =3 technical replicates and adjusted to OD_600_= 0.5.

In the inducible expression system (**Figures 4C and 4D**), changes in luminescence occurred more gradually following the addition of IPTG and antibiotics in both LuxCDABE and e*Cs*αDOX-eLuxA expressing cells. In *E. coli* cells expressing LuxCDABE, luminescence continued to increase for approximately 1 hour after IPTG induction in the presence of antibiotics but subsequently declined after ~5 hours and ~3 hours of exposure to meropenem and kanamycin, respectively. In contrast, following rifampicin treatment, LuxCDABE exhibited only a modest initial increase, followed by a pronounced decrease. Consistent with the results obtained using the constitutive expression system, cell expressing e*Cs*αDOX-eLuxA maintained sustained luminescence under all antibiotic conditions, with a particularly increase observed upon rifampicin treatment.

Consistent with previous reports^32^, these results indicate that shortly after induction and antibiotic treatment, *E. coli* JM109(DE3) cells retain sufficient metabolic capacity to support heterologous protein expression. During this early phase, host cells maintain adequate levels of ATP synthase and ATP-associated enzymes, allowing ATP-dependent LuxEC-mediated luciferin biosynthesis to proceed over a limited time scale. In addition, because rifampicin inhibits the native *E. coli* RNA polymerase but does not affect T7 RNA polymerase^32,33^, gene expression driven by the T7 system in both LuxCDABE and e*Cs*αDOX–eLuxA can be sustained transiently. However, in the LuxCDABE system, bioluminescence eventually declines, likely due to depletion of intracellular ATP or reduced activity of ATP-dependent enzymes required for luciferin biosynthesis^11^. In contrast, the ATP-independent e*Cs*αDOX-eLuxA system exhibited higher and more sustained luminescence, particularly under rifampicin treatment in the inducible expression system, compared with carbenicillin-treated controls. Notably, rifampicin has also been reported to enhance heterologous protein expression by increasing the availability or activity of molecular chaperones involved in recombinant protein folding^32^, which may further contribute to the enhanced luminescence observed under these conditions.

## DISCUSSION

In this study, we established an alternative autonomous bioluminescence pathway derived from the bacterial bioluminescence system that operates independently of ATP. In this pathway, α-dioxygenase (αDOX) catalyzes the conversion of fatty acids into fatty aldehydes without requiring ATP or NADPH, thereby functionally replacing the ATP-dependent LuxEC complex as the luciferin supply module for bacterial luciferase. αDOXs are widely distributed across the plant kingdom and are also found in non-plant organisms such as cyanobacteria. In plants, *Arabidopsis thaliana* αDOX (*At*αDOX) and *Oryza sativa* αDOX (*Os*αDOX) have been extensively characterized, with available crystal structures, and show a preference for longer-chain fatty acids (>C16)^20,34^. In contrast, αDOXs from cyanobacteria, including *Crocosphaera subtropica* (*Cs*αDOX), *Calothrix parietina* (*Cal*αDOX), and *Leptolyngbya* sp. (*Lep*αDOX) have been reported to preferentially utilize medium-chain fatty acids^15,22^. Given that medium-chain fatty acids are more compatible with the Lux pathway for fatty aldehyde production, these enzymes were selected with the expectation of enhancing luminescence in an ATP-independent manner.

Although the bioluminescence intensity of the *Cs*αDOX–Lux operon remained lower than that of the native LuxCDABE operon, this difference can be attributed to several factors. First, *Cs*αDOX produces (n−1) fatty aldehydes, and differences in preferred carbon chain length may reduce substrate compatibility with Lux luciferase^13^. Second, the catalytic activity of *Cs*αDOX *in vivo* was lower than that of the ATP-dependent LuxEC pathway (**Figure S2**), resulting in reduced fatty aldehyde production and, consequently, lower luminescence intensity^20^. As αDOX produces an n−1 fatty aldehyde, which may contribute to the reduced luminescence compared to the native LuxCDABE operon, alternative biosynthetic pathways for generating fatty aldehydes from fatty acids have been reported. These include fatty acyl-CoA/ACP reductases (FAR)^35^ and carboxylic acid reductases (CAR)^36^, both of which can produce fatty aldehydes with the same carbon chain length as the corresponding fatty acids. However, these enzymes require additional cofactors such as ATP and NADPH, and in the case of CAR, an additional partner enzyme, phosphopantetheinyltransferase (PPTase), for activation^36^. Therefore, while such pathways may restore aldehyde chain length, they would compromise the ATP-independent advantage of the αDOX-based system.

We further optimized the catalytic activity of *Cs*αDOX by combining two machine learning–guided mutagenesis approaches, EvoProtGrad^26^ and EVOLVEpro^27^. The models were trained to explore the protein fitness and activity landscape by predicting sequence–function relationships and prioritizing variants with improved target properties (i.e., bioluminescence output). Through iterative prediction and selection, the network identifies mutations that are expected to enhance enzyme performance while maintaining structural stability. The resulting mutations are predicted to improve protein fitness by modulating key properties such as folding efficiency, catalytic activity, and/or substrate accessibility. Further structural characterization, such as crystallographic analysis of the mutants, will help elucidate the underlying mechanisms responsible for the observed improvements. This combined approach significantly increased the luminescence output of the ATP-independent Lux system, ultimately enabling its detection by single-cell bioluminescence imaging.

The successful imaging of the ATP-independent autonomous bioluminescence system using the e*Cs*αDOX–eYNLX operon provides a valuable alternative tool for monitoring biological phenomena under metabolic perturbation in living bacteria. Although further optimization is still required, this system currently requires longer exposure times (approximately 3 min) compared to ATP-dependent system using LuxCDABE operon. This longer exposure time may limit temporal resolution and dynamic imaging. However, for applications such as monitoring gene expression, which typically occur on minute-to-hour timescales, the current performance is sufficient to generate reliable signal responses. Moreover, when combined with Lux color variants with NLXs system^12^, this platform offers strong potential for multiplexed imaging of multiple biological processes under varying metabolic conditions in living cells.

Finally, we investigated whether the ATP-independent Lux system responds to antibiotic treatment in the same manner as the native LuxCDABE operon. Because antibiotics can strongly affect ATP production and overall metabolic activity in bacteria^37,38^. treatment with antibiotics—including a translation inhibitor (kanamycin), a cell wall biosynthesis inhibitor (meropenem), and a transcription inhibitor (rifampicin) —led to a marked decrease in bioluminescence intensity from the LuxCDABE operon. In contrast, this decrease was not observed in the ATP-independent Lux system. Moreover, under inducible promoter control in *E. coli* JM109(DE3), rifampicin strongly suppressed luminescence from LuxCDABE, whereas the ATP-independent Lux system exhibited an increase in luminescence following inducer addition and rifampicin treatment.

Antibiotics such as kanamycin, meropenem, and rifampicin affect multiple aspects of cellular physiology, including transcription, translation, and overall metabolic activity. Therefore, the observed differences in luminescence cannot be attributed solely to ATP availability ^39,40^. While the ATP-independent nature of the αDOX–Lux–based system may contribute to its relative robustness, it is likely that a combination of factors—including broader metabolic perturbations—also influence the observed signal output. Further studies using more controlled perturbations of cellular energy metabolism will be necessary to directly assess the specific contribution of ATP dependence to αDOX-based system performance. For example, previous work using minimal *E. coli* cells (SimCells) has shown that Lux-based luminescence is reduced in the absence of glycolysis-encoding plasmids compared to SimCells expressing glycolysis pathways, highlighting the importance of cellular energy metabolism for signal output ^41^. In this context, ATP-independent systems such as αDOX–Lux may provide a useful platform for probing metabolic states in simplified or energy-limited cellular systems and could be further developed as sensors for studying minimal cells.

Bacterial bioluminescence systems require ATP and NADPH to produce fatty aldehydes. LuxE and LuxC form a complex that produces fatty aldehydes in an ATP- and NADPH-dependent manner. In contrast, *Cs*αDOX utilizes fatty acids (n−1) as substrates without requiring ATP or NADPH. Nevertheless, both systems rely on the intracellular availability of fatty acids to generate detectable bioluminescence signals, as shown in **Figure S2**. In bacteria, fatty acids are synthesized via the type II fatty acid synthase (FASII) pathway, which uses malonyl-CoA as the initial substrate for elongation through fatty acid synthase (Fab) enzymes, producing fatty acyl-ACP intermediates^42,43^. Although the elongation cycle itself does not require ATP, the formation of malonyl-CoA from acetyl-CoA by acetyl-CoA carboxylase is ATP-dependent^42^. Importantly, the Lux pathway does not directly incorporate malonyl-CoA as an extender unit in its luminescence pathway, in contrast to the fungal Luz pathway, where hispidin is synthesized from malonyl-CoA as extender compound by HispS (**Figure S1**). In addition to fatty aldehyde biosynthesis, NADH is also required for the generation of FMNH_2_ from FMN via native flavin reductases, such as Fre, in *E. coli*. Nevertheless, expression of the luxCDABE operon without exogenous Fre in *E. coli* is sufficient to produce bioluminescence signals above background levels, a phenomenon also observed in the ATP-independent Lux system developed here. Notably, previous studies have shown that deletion of *fre* in *E. coli* does not significantly reduce luminescence intensity relative to wild-type strain^11,13^. These findings suggest that FMNH_2_ production is supported by multiple endogenous redox enzymes rather than relying exclusively on Fre, and that redundant or cooperative redox pathways may contribute to flavin reduction—an aspect that warrants further investigation. Overall, the ATP-free fatty aldehyde biosynthesis Lux system, constructed by combining enhanced *Cs*αDOX (e*Cs*αDOX) with enhanced LuxA (eLuxA), provides a metabolically robust platform for monitoring biological phenomena in bacteria with reduced sensitivity to metabolic fluctuations.

### Limitations of the study

The antibiotic-based experiments presented in this study should be interpreted with caution due to inherent limitations in their physiological relevance. In particular, the use of a T7-driven expression system in combination with rifampicin creates a non-physiological condition in which host transcription is largely inhibited while heterologous expression is maintained. This setup may amplify differences between the αDOX–Lux and LuxCDABE systems and does not fully reflect typical cellular environments. Moreover, the observed differences in luminescence under antibiotic treatment cannot be attributed solely to ATP availability, but rather likely arise from a combination of overlapping factors. While the αDOX–Lux system shows potential under energy-limited or perturbed conditions, its current performance remains constrained for certain applications, such as promoter-based biosensing, and further optimization will be required. In addition, this system is its dependence on molecular oxygen, as both LuxA and *Cs*αDOX require oxygen for catalytic activity. Consequently, further protein and pathway engineering will be necessary to extend the applicability of this system to anaerobic or microaerophilic bacteria, potentially through the development of oxygen-tolerant enzyme variants.

## RESOURCE AVAILABILITY

### Lead contact

Further information and requests for resources and reagents should be directed to and will be fulfilled by the lead contact, Takeharu Nagai.

### Materials availability

Plasmids generated in this study are available from the authors on reasonable request.

### Data and code availability

- Data reported in this paper will be shared by the lead contact upon request.
- This paper does not report original code.
- Any additional information required for data reported in this paper is available from the lead contact.

## ACKNOWLEDGMENTS

This work was financially supported by grants from the New Energy and Industrial Technology Development Organization (No. 22681865 to T.N).

## AUTHOR CONTRIBUTIONS

Conceptualization, S.H.K. and T.N.; methodology, S.H.K.; Investigation, S.H.K.; writing—original draft, S.H.K.; writing—review & editing, S.H.K. and T.N.; funding acquisition, T.N.; resources, S.H.K. and T.N.; supervision, S.H.K. and T.N.

## SUPPLEMENTAL INFORMATION

**Document S1. Figures S1–S5 and Tables S1**

## METHODS

### KEY RESOURCES TABLE

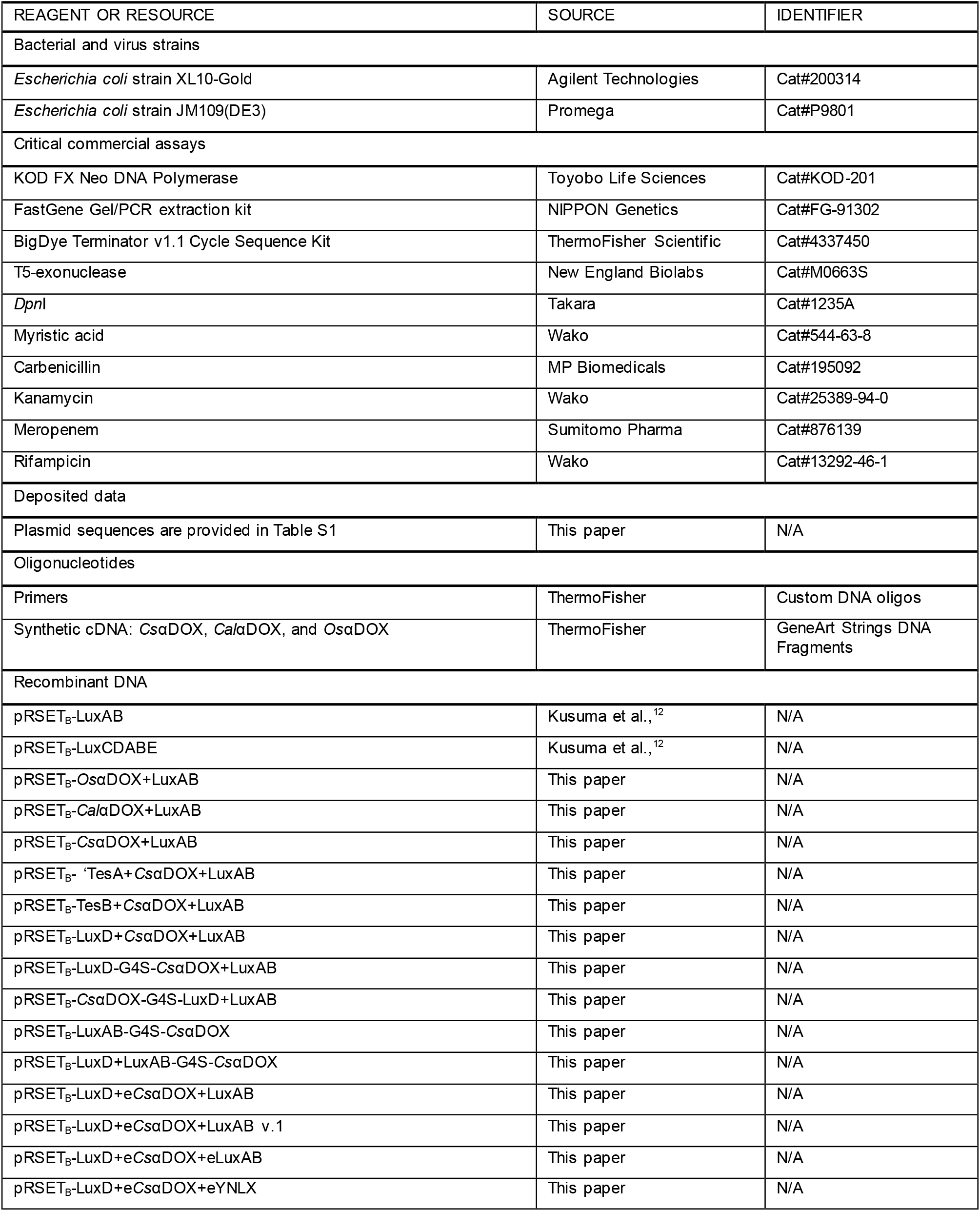
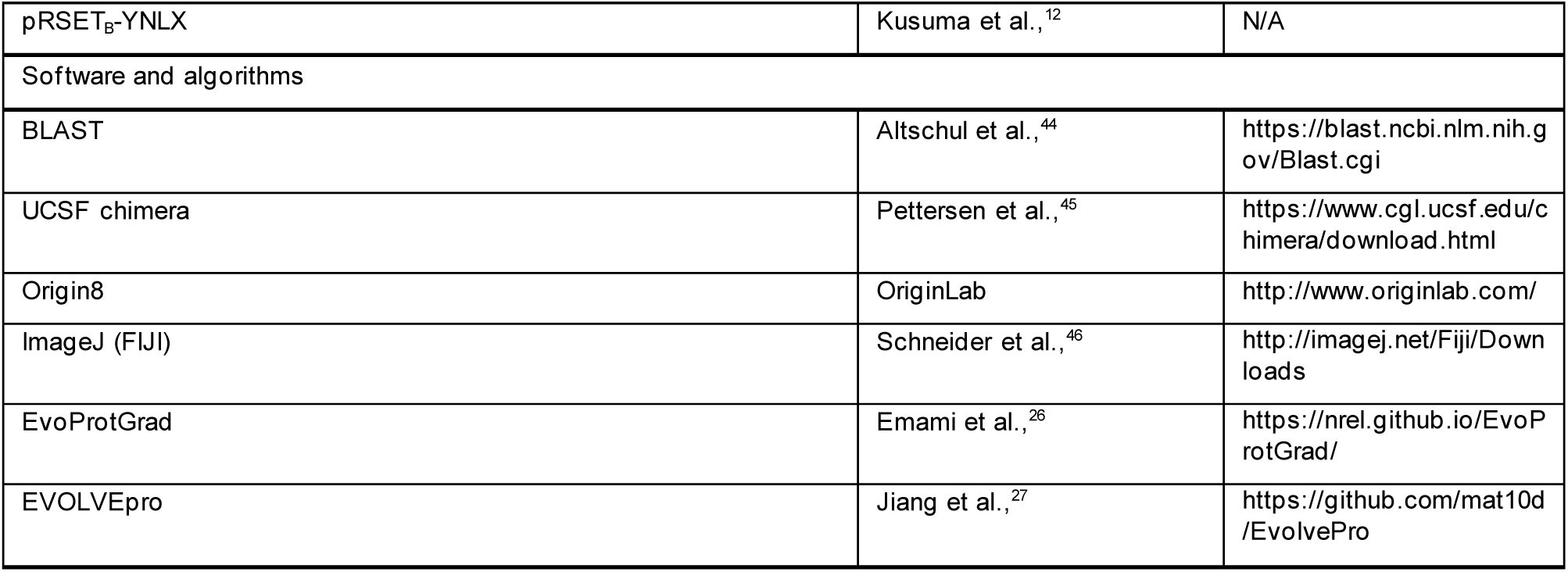

## METHOD DETAILS

### General methods

DNA oligonucleotides were purchased from ThermoFisher Scientific. KOD-FX-Neo (Toyobo Life Science) was used for PCR amplification. PCR products were purified by agarose gel electrophoresis using a FastGene Gel/PCR extraction kit (NIPPON Genetics). DpnI-treated was used to degrade the parental vector from the inverse-PCR vector. Small-scale plasmid DNA was obtained from a 1.5 mL LB-liquid bacterial culture by alkaline lysis and ethanol precipitation. DNA sequencing of the cDNA constructs was performed using the BigDye Terminator v1.1 Cycle Sequencing kit (Life Technologies).

### Construction of αDOX-Lux

The luxAB operon in the pRSET_B_ vector was used as the starting material for constructing the ATP-independent Lux system in *E. coli. E. coli* codon-optimized sequences of *Os*αDOX, *Cal*αDOX, and *Cs*αDOX were synthesized by ThermoFisher Scientific. The cDNAs of all αDOX genes were amplified by PCR and subcloned into LuxAB operon to yield αDOX-LuxAB/pRSET_B_ using T5 exonuclease–mediated low-temperature DNA cloning method (TLTC) ^47^. For thiosterases cloning, ‘TesA and TesB were amplified from *E. coli* K-12 genome, and LuxD was amplified from LuxCDABE/pRSET_B_ vector. PCR products were subcloned into *Cs*αDOX-LuxAB/pRSET_B_ to yield thiosterases-*Cs*αDOX-LuxAB/pRSET_B_ using TLTC assembly. For fusion strategy, LuxD was fused wot either the N- or C-terminus of *Cs*αDOX using a GGGGS linker, and *Cs*αDOX was fused to the C-terminus of LuxB using the same linker to generate LuxB-GGGGS-CsαDOX, assembled by TLTC method. Optimized variants of *Cs*αDOX and LuxA obtained by mutagenesis were subcloned into the LuxD-*Cs*αDOX-LuxAB operon. Venus was fused to eLuxA with EL linker by TLTC assembly.

### Machine learning–guided directed evolution of *Cs*αDOX and LuxA

Mutagenesis of *Cs*αDOX and LuxA guided by two approaches of machine learning, EvoProtGrad^26^ and EVOLVEpro^27^. For EvoProtGrad, the source code was available at https://nrel.github.io/EvoProtGrad/. For EVOLVEpro, the source code was available at https://github.com/mat10d/EvolvePro. Both of models used ESM2-3B base layer protein language model (PLM) for evolution. The wild-type sequences of LuxA and *Cs*αDOX were initially used as inputs for the EvoProtGrad framework, employing two expert models consisting of a protein language model (PLM) and a CNN model trained on GFP-related fitness datasets. The Markov chain Monte Carlo (MCMC) parameter was set to 100 iterations, with a maximum of 12 mutations per variant. Candidate variants predicted were selected for the next optimization round through EVOLVEpro. The resulting candidates were experimentally evaluated for luminescence activity by introducing mutations using site-directed mutagenesis (SDM). Variants from each screening round were subsequently used as inputs for the next round of EVOLVEpro-based activity optimization, generating up to 12 new mutations per cycle through round 4. For multimutant construction, we manually selected at least the top three best-performing variants from each round and combined beneficial mutations to generate enhanced LuxA and CsαDOX variants. Site-directed mutagenesis (SDM) with overlapping primers was performed to obtain the mutated variants using LuxD-*Cs*αDOX-LuxAB/pRSET_B_ and LuxCDABE/pRSET_B_ as template. The following thermal cycling conditions were used for SDM: 18 cycles of denaturation (94° C, 15 s), annealing (55° C, 30 s), and extension (68° C, 5 min). The PCR products were treated by *Dpn*I and transformed into *E. coli* JM109(DE3).

### Luciferase assay

The modified plasmids were transformed into *E. coli* JM109(DE3) and expressed constitutively under the T7 promoter. Cultures were grown overnight, plated in 96-well white plates (Greiner Bio-One, 150 μL/well), and bioluminescence intensity was measured using a multimode plate reader (Spectra Max iD5, Molecular Devices) for 3 s of exposure. Bioluminescence values (including mutagenesis) were normalized to OD_600_ to obtain the final bioluminescence intensity. For antibiotic treatment experiments with the *eCs*αDOX-eLux operon and LuxCDABE, overnight cultures were adjusted to OD_600_ = 0.5 and bioluminescence was measured using the same plate reader for 3 s of exposure. The final concentrations of antibiotics were meropenem (30 µg/mL), kanamycin (50 µg/mL), rifampicin (25 µg/mL), and carbenicillin (100 µg/mL). IPTG was added to a final concentration of 1 mM. Several myristic acid (Wako) concentrations were added to a final concentration from 20 to 1000 µM.

### Bioluminescence imaging

Autonomous bioluminescence images of recombinant *E. coli* were picked into 10 µL of H_2_O. 0.3 µL of each recombinant *E. coli* was introduced into 35 mm glass bottom dishes using the agar pad method. Luminescence imaging was acquired with an inverted microscope based on the IXploreTM Live for Luminescence (EVIDENT) system equipped with an EM-CCD camera (Andor iXon Ultra 888) with several min of exposure, EM-gain of 1000×, ×60 objective lens and 2 × 2 binning settings. All imaging conditions were performed at 37 °C in a stage-top incubator, STX (TOKAI HIT). All bioluminescent images were analyzed using ImageJ software^46^. Background signals were subtracted from the raw images, and cosmic ray artifacts were removed using an adaptive median filter in ImageJ without altering the original brightness or cellular morphology. Pseudocolor rendering was applied to microscopy images, with the “Red Hot” lookup table used to represent total luminescence intensity.

### QUANTIFICATION AND STATISTICAL ANALYSIS

Data fitting and statistical analyses were analyzed using Origin8 software (OriginLab) or Microsoft Excel. Statistical values including the exact n and statistical significance are reported in Figure legends. Comparisons between two groups were performed using an unpaired Student’s *t-test*, while multiple group comparisons were analyzed by one-way ANOVA followed by post hoc Tukey’s honestly significant difference test.

## REFERENCES

1. Yuan, Z., Jiang, Q., and Liang, G. (2025). Inspired by nature: Bioluminescent systems for bioimaging applications. Talanta 281, 126821.

2. Mezzanotte, L., van ‘t Root, M., Karatas, H., Goun, E.A., and Löwik, C.W.G.M. (2017). In Vivo Molecular Bioluminescence Imaging: New Tools and Applications. Trends in Biotechnology 35, 640–652.

3. Kusuma, S.H., Hattori, M., and Nagai, T. (2025). Autonomous Bioluminescence Systems: From Molecular Mechanisms to Emerging Applications. JACS Au 5, 5237–5252.

4. Shimomura, O., and Yampolsky, I. V. (2019). Bioluminescence: Chemical principles and methods (3rd Edition).

5. Hall, M.P., Unch, J., Binkowski, B.F., Valley, M.P., Butler, B.L., Wood, M.G., Otto, P., Zimmerman, K., Vidugiris, G., MacHleidt, T., et al. (2012). Engineered luciferase reporter from a deep sea shrimp utilizing a novel imidazopyrazinone substrate. ACS Chem. Biol. 7. 1848–1857.

6. Nakajima, Y., Yamazaki, T., Nishii, S., Noguchi, T., Hoshino, H., Niwa, K., Viviani, V.R., and Ohmiya, Y. (2010). Enhanced beetle luciferase for high-resolution bioluminescence imaging. PLoS One 5.

7. Suzuki, K., Kimura, T., Shinoda, H., Bai, G., Daniels, M.J., Arai, Y., Nakano, M., and Nagai, T. (2016). Five colour variants of bright luminescent protein for real-time multicolour bioimaging. Nat. Commun. 7, e10011.

8. Su, Y., Walker, J.R., Hall, M.P., Klein, M.A., Wu, X., Encell, L.P., Casey, K.M., Liu, L.X., Hong, G., Lin, M.Z., et al. (2023). An optimized bioluminescent substrate for non-invasive imaging in the brain. Nat. Chem. Biol. 19, 731–739.

9. Tantama, M., Min, S.H., French, A.R., Trull, K.J., Tat, K., and Varney, S.A. (2019). Ratiometric BRET measurements of ATP with a genetically-encoded luminescent sensor. Sensors 19, 3502.

10. Kotlobay, A.A., Sarkisyan, K.S., Mokrushina, Y.A., Marcet-Houben, M., Serebrovskaya, E.O., Markina, N.M., Somermeyer, L.G., Gorokhovatsky, A.Y., Vvedensky, A., Purtov, K. V., et al. (2018). Genetically encodable bioluminescent system from fungi. Proc. Natl. Acad. Sci. U. S. A. 115, 12728–12732.

11. Gregor, C., Gwosch, K.C., Sahl, S.J., and Hell, S.W. (2018). Strongly enhanced bacterial bioluminescence with the ilux operon for single-cell imaging. Proc. Natl. Acad. Sci. U. S. A. 115, 962–967.

12. Kusuma, S.H., Kakizuka, T., Hattori, M., and Nagai, T. (2024). Autonomous multicolor bioluminescence imaging in bacteria, mammalian, and plant hosts. Proc. Natl. Acad. Sci. USA 121, e2406358121.

13. Gregor, C. (2022). Generation of bright autobioluminescent bacteria by chromosomal integration of the improved lux operon ilux2. Sci. Rep. 12, 19039.

14. Bozcal, E., Dagdeviren, M., Uzel, A., and Skurnik, M. (2017). LuxCDE-luxAB-based promoter reporter system to monitor the Yersinia enterocolitica O:3 gene expression in vivo. PLoS One 12, e0172877.

15. Hammer, A.K., Albrecht, F., Hahne, F., Jordan, P., Fraatz, M.A., Ley, J., Geissler, T., Schrader, J., Zorn, H., and Buchhaupt, M. (2020). Biotechnological Production of Odor-Active Methyl-Branched Aldehydes by a Novel α-Dioxygenase from Crocosphaera subtropica. J. Agric. Food Chem. 68, 10432–10440.

16. Hamberg, M., Ponce de Leon, I., Rodriguez, M.J., and Castresana, C. (2005). α-Dioxygenases. Biochem. Biophys. Res. Commun. 338, 169–174.

17. Kim, I.J., Bayer, T., Terholsen, H., and Bornscheuer, U.T. (2022). α-Dioxygenases (α-DOXs): Promising Biocatalysts for the Environmentally Friendly Production of Aroma Compounds. ChemBioChem 23, e202100693.

18. Meighen, E.A. (1991). Molecular biology of bacterial bioluminescence. Microbiol. Rev. 55, 123–142.

19. Tian, Q., Wu, J., Xu, H., Hu, Z., Huo, Y., and Wang, L. (2022). Cryo-EM structure of the fatty acid reductase LuxC–LuxE complex provides insights into bacterial bioluminescence. J. Biol. Chem. 298, 102006.

20. Zhu, G., Koszelak-Rosenblum, M., and Malkowski, M.G. (2013). Crystal structures of α-dioxygenase from Oryza sativa: Insights into substrate binding and activation by hydrogen peroxide. Protein Science 22, 1432–1438.

21. Kaehne, F., Buchhaupt, M., and Schrader, J. (2011). A recombinant α-dioxygenase from rice to produce fatty aldehydes using E. coli. Appl. Microbiol. Biotechnol. 90, 989–995.

22. Kim, I.J., Brack, Y., Bayer, T., and Bornscheuer, U.T. (2022). Two novel cyanobacterial α-dioxygenases for the biosynthesis of fatty aldehydes. Appl. Microbiol. Biotechnol. 106, 197–210.

23. Cho, H., and Cronan, J.E. (1995). Defective export of a periplasmic enzyme disrupts regulation of fatty acid synthesis. J. Biol. Chem. 270, 4216–4219.

24. Steen, E.J., Kang, Y., Bokinsky, G., Hu, Z., Schirmer, A., McClure, A., Del Cardayre, S.B., and Keasling, J.D. (2010). Microbial production of fatty-acid-derived fuels and chemicals from plant biomass. Nature 463, 559–562.

25. Li, J., Derewenda, U., Dauter, Z., Smith, S., and Derewenda, Z.S. (2000). Crystal structure of the Escherichia coli thioesterase II, a homolog of the human Nef binding enzyme. Nat. Struct. Biol. 7. 555–559.

26. Emami, P., Perreault, A., Law, J., Biagioni, D., and St. John, P. (2023). Plug & play directed evolution of proteins with gradient-based discrete MCMC. Mach. Learn. Sci. Technol. 4, 025014.

27. Jiang, K., Yan, Z., Bernardo M. Di, Sgrizzi, S.R., Villiger, L., Kayabolen, A., Kim, B.J., Carscadden, J.K., Hiraizumi, M., Nishimasu, H., et al. (2025). Rapid in silico directed evolution by a protein language model with EVOLVEpro. Science 387, eadr6006.

28. Lin, Z., Akin, H., Rao, R., Hie, B., Zhu, Z., Lu, W., Smetanin, N., Verkuil, R., Kabeli, O., Shmueli, Y., et al. (2023). Evolutionary-scale prediction of atomic-level protein structure with a language model. Science 379, 1123–1130.

29. Krause, K.M., Serio, A.W., Kane, T.R., and Connolly, L.E. (2016). Aminoglycosides: An overview. Cold Spring Harb. Perspect. Med. 6, a027029.

30. Zandi, T.A., and Townsend, C.A. (2021). Competing off-loading mechanisms of meropenem from an L,D-transpeptidase reduce antibiotic effectiveness. Proc. Natl. Acad. Sci. USA 118.

31. Campbell, E.A., Korzheva, N., Mustaev, A., Murakami, K., Nair, S., Goldfarb, A., and Darst, S.A. (2001). Structural mechanism for rifampicin inhibition of bacterial RNA polymerase. Cell 104. 901–912.

32. Kuderová, A., Nanak, E., Truksa, M., and Brzobohatý, B. (1999). Use of rifampicin in T7 RNA polymerase-driven expression of a plant enzyme: Rifampicin improves yield and assembly. Protein Expr. Purif. 16, 405–409.

33. Tabor, S., and Richardson, C.C. (1985). A bacteriophage T7 RNA polymerase/promoter system for controlled exclusive expression of specific genes. Proc. Natl. Acad. Sci. USA 82, 1074–1078.

34. Goulah, C.C., Zhu, G., Koszelak-Rosenblum, M., and Malkowski, M.G. (2013). The crystal structure of α-dioxygenase provides insight into diversity in the cyclooxygenase-peroxidase superfamily. Biochemistry 52. 10.1021/bi400013k.

35. Foo, J.L., Rasouliha, B.H., Susanto, A.V., Leong, S.S.J., and Chang, M.W. (2020). Engineering an Alcohol-Forming Fatty Acyl-CoA Reductase for Aldehyde and Hydrocarbon Biosynthesis in Saccharomyces cerevisiae. Front. Bioeng. Biotechnol. 8. 10.3389/fbioe.2020.585935.

36. Kalim Akhtara, M., Turner, N.J., and Jones, P.R. (2013). Carboxylic acid reductase is a versatile enzyme for the conversion of fatty acids into fuels and chemical commodities. Proc. Natl. Acad. Sci. U. S. A. 110. 10.1073/pnas.1216516110.

37. Stokes, J.M., Lopatkin, A.J., Lobritz, M.A., and Collins, J.J. (2019). Bacterial Metabolism and Antibiotic Efficacy. Cell Metab. 30, 251–259.

38. Maglica, Ž., Özdemir, E., and McKinney, J.D. (2015). Single-cell tracking reveals antibiotic-induced changes in mycobacterial energy metabolism. mBio 6.

39. Lobritz, M.A., Belenky, P., Porter, C.B.M., Gutierrez, A., Yang, J.H., Schwarz, E.G., Dwyer, D.J., Khalil, A.S., and Collins, J.J. (2015). Antibiotic efficacy is linked to bacterial cellular respiration. Proc. Natl. Acad. Sci. U. S. A. 112.

40. Li, B., Srivastava, S., Shaikh, M., Mereddy, G., Garcia, M.R., Chiles, E.N., Shah, A., Ofori-Anyinam, B., Chu, T.Y., Cheney, N.J., et al. (2025). Bioenergetic stress potentiates antimicrobial resistance and persistence. Nat. Commun. 16.

41. Fan, C., Davison, P.A., Habgood, R., Zeng, H., Decker, C.M., Salazar, M.G., Lueangwattanapong, K., Townley, H.E., Yang, A., Thompson, I.P., et al. (2020). Chromosome-free bacterial cells are safe and programmable platforms for synthetic biology. Proc. Natl. Acad. Sci. U. S. A. 117.

42. Janßen, H.J., and Steinbüchel, A. (2014). Fatty acid synthesis in Escherichia coli and its applications towards the production of fatty acid based biofuels. Biotechnol. Biofuels 7.

43. Yu, X., Liu, T., Zhu, F., and Khosla, C. (2011). In vitro reconstitution and steady-state analysis of the fatty acid synthase from Escherichia coli. Proc. Natl. Acad. Sci. U. S. A. 108.

44. Altschul, S.F., Madden, T.L., Schäffer, A.A., Zhang, J., Zhang, Z., Miller, W., and Lipman, D.J. (1997). Gapped BLAST and PSI-BLAST: A new generation of protein database search programs. Nucleic Acids Research 25, 3389–3402.

45. Pettersen, E.F., Goddard, T.D., Huang, C.C., Couch, G.S., Greenblatt, D.M., Meng, E.C., and Ferrin, T.E. (2004). UCSF Chimera - A visualization system for exploratory research and analysis. J. Comput. Chem. 25.

46. Schneider, C.A., Rasband, W.S., and Eliceiri, K.W. (2012). NIH Image to ImageJ: 25 years of image analysis. Nat. Methods 9, 671–675.

47. Yu, F., Li, X., Wang, F., Liu, Y., Zhai, C., Li, W., Ma, L., and Chen, W. (2023). TLTC, a T5 exonuclease–mediated low-temperature DNA cloning method. Front. Bioeng. Biotechnol. 11.

