## Supplemental Information for "ATP-Free Fatty Aldehyde Biosynthesis Enables an Autonomous Lux-Based Bioluminescence System"

### TABLE OF CONTENTS

Figures S1-S5

Table S1

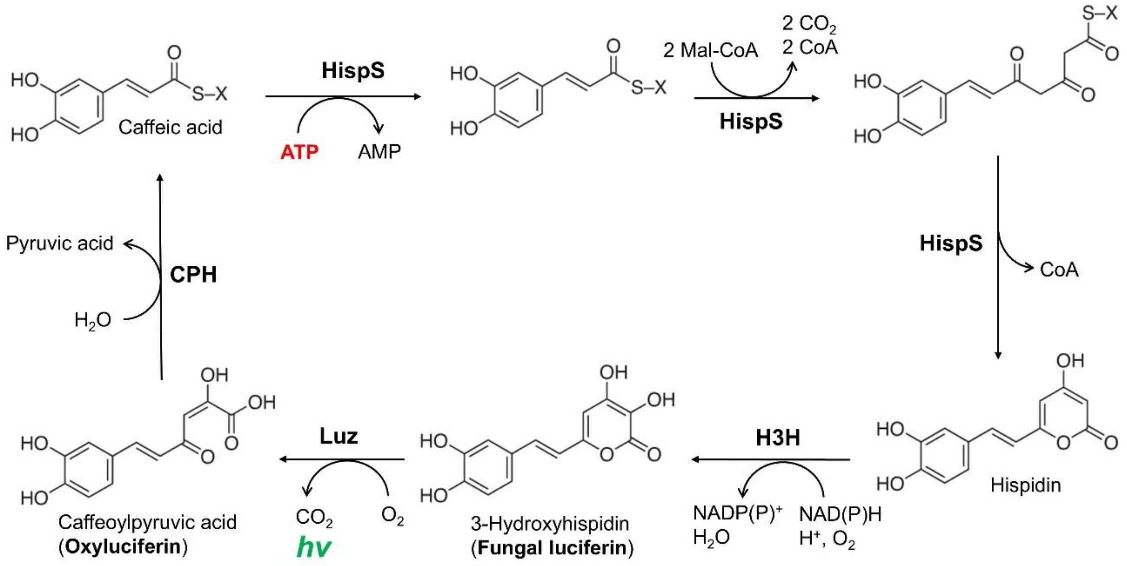

**Figure S1.** Fungal bioluminescence pathway.

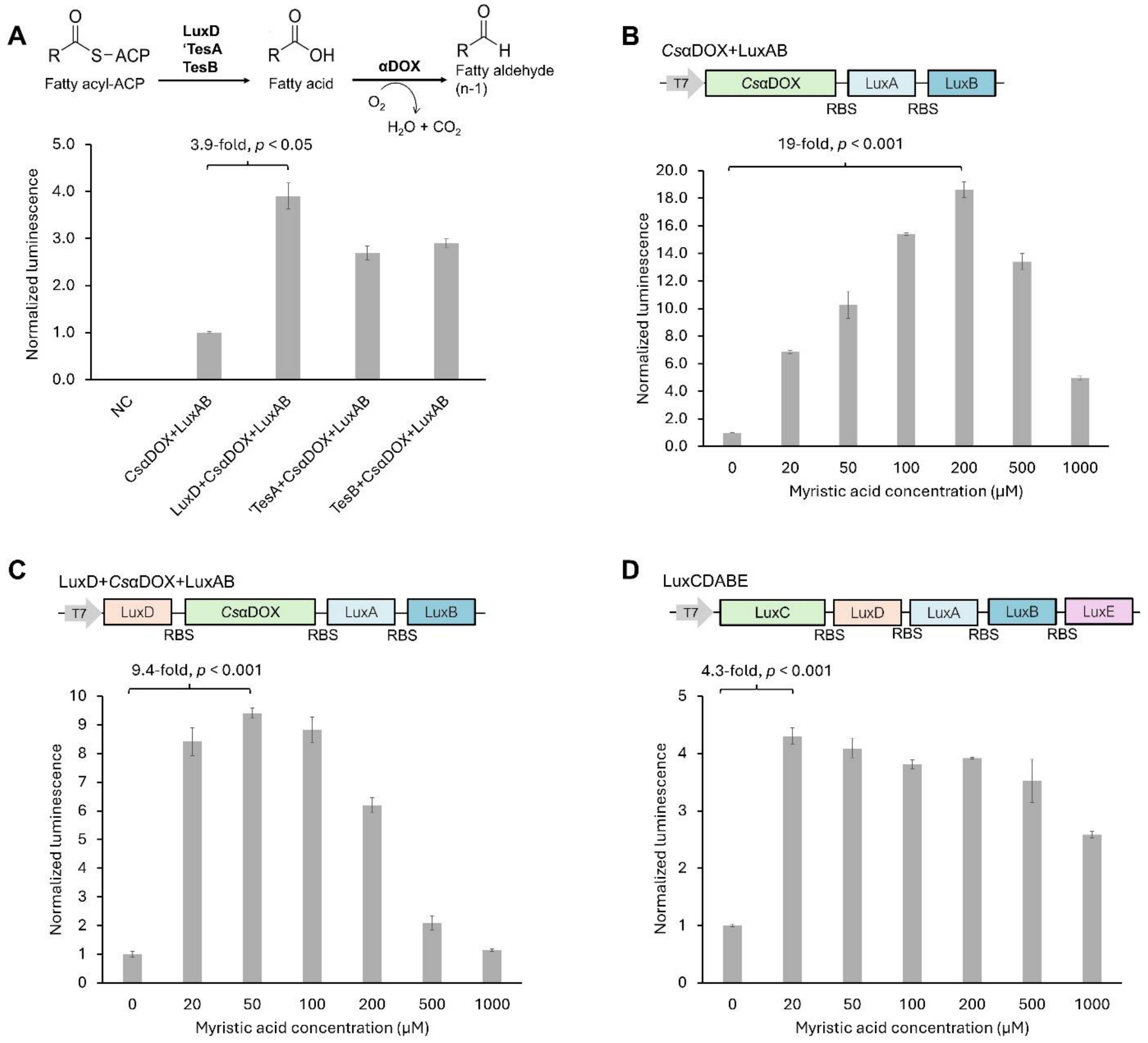

**Figure S2. Optimization of thiosterases to *Cs*αDOX-LuxAB**. (A) Schematic of the thioesterase-mediated fatty acid production pathway (upper panel) and bioluminescence intensity of thioesterase variants in the *Cs*αDOX-Lux operon (bottom). NC, non-transformant *E. coli* JM109(DE3). Data are presented as mean ± SD. *n* =3 technical replicates and normalized by $OD_{600}$. Bioluminescence intensity of *E. coli* expressing *Cs*αDOX+LuxAB (B), LuxD+*Cs*αDOX+LuxAB (C), and LuxCDABE (D) with $OD_{600}$= 0.5, upon addition of several concentrations of myristic acid. Data are presented as mean ± SD. *n* =3 technical replicates.

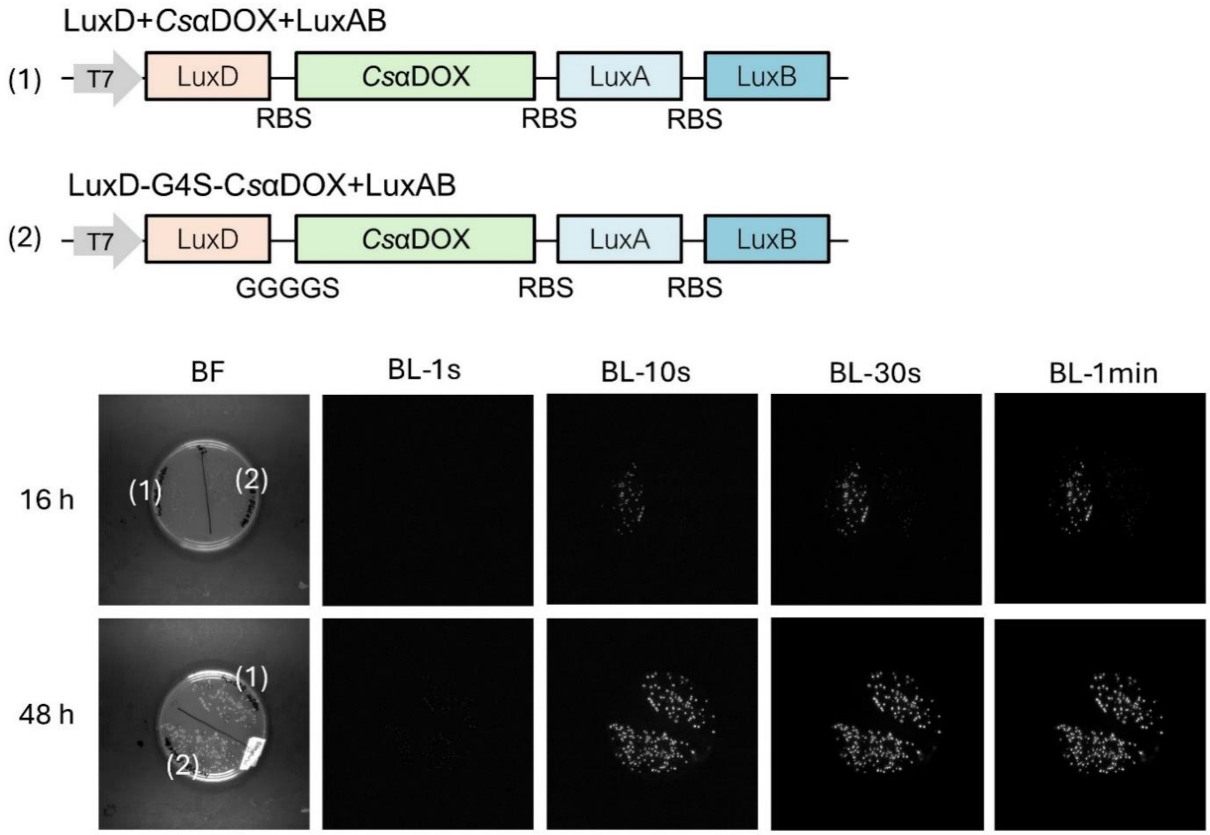

**Figure S3. Fusion strategy to *Cs*αDOX-LuxAB**. Schematic of the fusion strategy linking LuxD to *Cs*αDOX-LuxAB (upper panel) and corresponding bioluminescent colony images after 16 h and 48 h of incubation (bottom). BF and BL are brightfield and bioluminescence intensity with several exposure time, respectively.

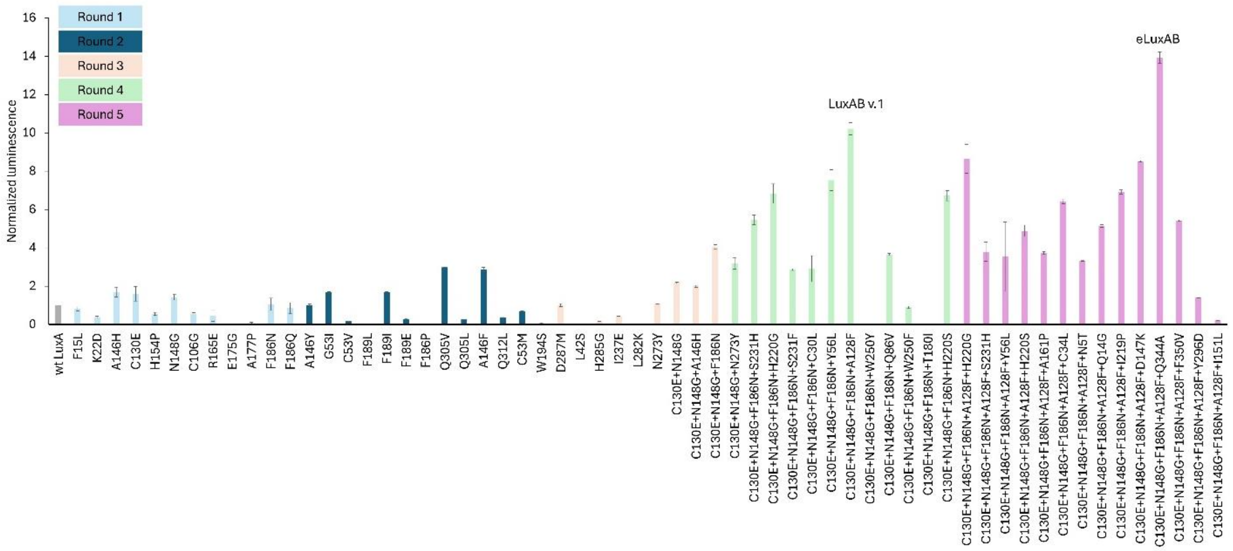

**Figure S4.** Screening of LuxA libraries by machine learning–guided directed evolution (EvoProtGrad and EVOLVEpro). Data are presented as mean ± SD. *n* =3 technical replicates and normalized by $OD_{600}$.

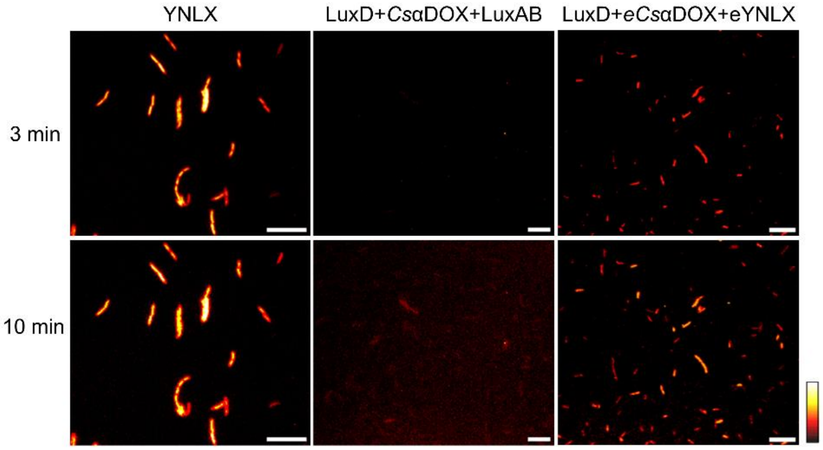

**Figure S5.** Single-cell bioluminescence imaging of *E. coli* expressing YNLX, *Cs*αDOX-Lux, and e*Cs*αDOX-eYNLX operon. Pseudocolor images, scale bars, 20μm; 60× magnification with indicated exposure times.

**Table S1**. Plasmids used in this study

| No | ID | Plasmid description | Plasmid map | Note |
|---|---|---|---|---|
| 1 | LuxAB | LuxA-RBS-LuxB/pRSET$_{B}$ | https://benchling.com/s/seq-OEurvDJa5Gs07PCShORq?m=slm-s7lEVClaTRXQl26ziBe8 | |
| 2 | LuxCDABE | LuxC-RBS-LuxD-RBS-LuxA-RBS-LuxB-RBS-LuxE/pRSET$_{B}$ | https://benchling.com/s/seq-CTNT3SXQNARnotBX5n1k?m=slm-d6RBAmvvtgy24rZ0DkFV | |
| 3 | *Os*αDOX+LuxAB | *Os*αDOX-RBS-LuxA-RBS-LuxB/pRSET$_{B}$ | https://benchling.com/s/seq-DP5IlcRHKxEv6VmUSLNN?m=slm-Td3Jg2ZkAE7YaRzygMqf | |
| 4 | *Cal*αDOX+LuxAB | *Cal*αDOX-RBS-LuxA-RBS-LuxB/pRSET$_{B}$ | https://benchling.com/s/seq-nXYkNvzPKVPknGEc72WV?m=slm-W9lDNCjlWBpIFT9L8ljJ | |
| 5 | *Cs*αDOX+LuxAB | *Cs*αDOX-RBS-LuxA-RBS-LuxB/pRSET$_{B}$ | https://benchling.com/s/seq-GiAkGkJDzHG7Q3mtdVyD?m=slm-gwdc8qkNBvk8UcJ48FjZ | |
| 6 | ‘TesA+*Cs*αDOX+LuxAB | ‘TesA-RBS-*Cs*αDOX-RBS-LuxA-RBS-LuxB/pRSET$_{B}$ | https://benchling.com/s/seq-ZJOe0svKH59kLgq6eSKx?m=slm-lTuLfQEPgbm2Tv1eRkUO | |
| 7 | TesB+*Cs*αDOX+LuxAB | TesB-RBS-*Cs*αDOX-RBS-LuxA-RBS-LuxB/pRSET$_{B}$ | https://benchling.com/s/seq-1UzXRSyH0UjgAC8FKAmV?m=slm-eOwiuwprJtDO4xrpq5XU | |
| 8 | LuxD+*Cs*αDOX+LuxAB | LuxD-RBS-*Cs*αDOX-RBS-LuxA-RBS-LuxB/pRSET$_{B}$ | https://benchling.com/s/seq-W7nQlPzqWCWdU1GnZEq1?m=slm-1bxb2jGZXAEV6mMGLXYq | |
| 9 | LuxD-G4S-*Cs*αDOX+LuxAB | LuxD-GGGGS-*Cs*αDOX-RBS-LuxA-RBS-LuxB/pRSET$_{B}$ | https://benchling.com/s/seq-KjP9GQQ54TGiNWri3G3b?m=slm-RzTGeRyDk5sSljPKW936 | |
| 10 | *Cs*αDOX-G4S-LuxD+LuxAB | *Cs*αDOX-GGGGS-LuxD-RBS-LuxA-RBS-LuxB/pRSET$_{B}$ | https://benchling.com/s/seq-aFiWsG5O5trtrPY45ENW?m=slm-1h1icpiynul0OgTFI45H | |
| 11 | LuxAB-G4S-*Cs*αDOX | LuxA-RBS-LuxB-GGGGS-*Cs*αDOX /pRSET$_{B}$ | https://benchling.com/s/seq-5e0mHcVhsuJ22YbeNL9w?m=slm-CS6zUWpK3HL7zn0hJ3ED | |
| 12 | LuxD+LuxAB-G4S-*Cs*αDOX | LuxD-RBS-LuxA-RBS-LuxB-GGGGS-*Cs*αDOX /pRSET$_{B}$ | https://benchling.com/s/seq-7WFRyumzaAgsUjKKMYZU?m=slm-YT7MDjvUWrxdcThPLqSB | |
| 13 | LuxD+e*Cs*αDOX+LuxAB | LuxD-RBS-e*Cs*αDOX-RBS-LuxA-RBS-LuxB/pRSET$_{B}$ | https://benchling.com/s/seq-aDcVnRCPzEDB9rDqM9fM?m=slm-3fAuVvPa57RkzZ7VCP4P | |
| 14 | LuxD+e*Cs*αDOX+LuxAB v.1 | LuxD-RBS-e*Cs*αDOX-RBS-LuxA v.1-RBS-LuxB/pRSET$_{B}$ | https://benchling.com/s/seq-XoXQWjZkE153kuBlB1bL?m=slm-4AcuxaFxleMLlESTQ1BD | |

| | | | |
|---|---|---|---|
| 15 | LuxD+e*Cs*αDOX+eLuxAB | LuxD-RBS-e*Cs*αDOX-RBS-eLuxA-RBS-LuxB/pRSET$_B$ | https://benchling.com/s/seq-NpyHhArvZ68qjsvhpwsj?m=slm-Mf7VkMIgRV7H9lVziGsT |
| 16 | LuxD+e*Cs*αDOX+eYNLX | LuxD-RBS-e*Cs*αDOX-RBS-VenusΔC10-EL-eLuxA-RBS-LuxB/pRSET$_B$ | https://benchling.com/s/seq-bBysTGxwgRmzqv4WfffI?m=slm-H7zpTo6ZrkNKI5F41BU0 |
